# Chromosomal rearrangements at hypomethylated Satellite 2 sequences are associated with impaired replication efficiency and increased fork stalling

**DOI:** 10.1101/554410

**Authors:** Yannick Delpu, Thomas McNamara, Patrick Griffin, Suhail Kaleem, Shubhada Narayan, Carl Schildkraut, Karen Miga, Mamta Tahiliani

**Affiliations:** Skirball Institute of Biomolecular Medicine/Department of Biochemistry and Molecular Pharmacology, New York University School of Medicine, New York City, NY, 10016, USA; Albert Einstein College of Medicine, Department of Cell Biology, New York, NY, 10461, USA; UC Santa Cruz Genomics Institute, University of California, Santa Cruz, California, USA

## Abstract

Cancer cells, aging cells, and cells from patients with the developmental disorder Immunodeficiency, <u>C</u>entromeric instability, and <u>F</u>acial anomalies (ICF) syndrome frequently display a striking loss of DNA methylation (hypomethylation) that is accompanied by increased DNA damage and chromosomal rearrangements. Despite the robust link, the mechanism by which hypomethylation leads to genomic instability is poorly understood. We report that the human pericentromeric repeat sequence Satellite 2 (SAT2) poses challenges to the DNA replication machinery when hypomethylated. Loss of methylation at SAT2 is associated with increased frequencies of chromosomal abnormalities and DNA damage. Hypomethylation of SAT2 is associated with elevated levels of replication stress signaling, and chromosomal abnormalities involving SAT2 are enhanced by low levels of aphidicolin-induced replication stress. To investigate the basis for these chromosomal abnormalities, we developed a single-molecule approach employing DNA combing to examine the progress of replication forks through SAT2 at the resolution of a single DNA molecule. Our analysis of replicating SAT2 molecules provides in vivo evidence that hypomethylation of SAT2 strongly decreases the efficiency of replicating these sequences suggesting that hypomethylation results in the formation of barriers to the replication machinery. Consistent with increased frequency of fork stalling at these sequences, we find increased levels of single-stranded DNA (ssDNA) binding protein RPA2 as well as asymmetric progression of sister replication forks within hypomethylated SAT2 sequences. Together these findings indicate that impaired replication triggers the formation of chromosomal aberrations observed at hypomethylated SAT2 sequences and also suggests a mechanistic basis for how the loss of DNA methylation may contribute to genomic instability in diverse pathological conditions.

## Introduction

Genome-wide hypomethylation is a frequent characteristic of tumor cells, aging cells, and cells from patients with the developmental disorder ICF syndrome. (2012; Cruickshanks et al., 2013; Ehrlich, 2002; Feinberg and Vogelstein, 1983) where it is associated with increased DNA damage and chromosomal rearrangements (Chen et al., 1998; Eden et al., 2003; Ehrlich et al., 2001; Gaudet et al., 2003; Hernandez et al., 1997). Despite the strong correction, the mechanism by which hypomethylation triggers genomic instability is poorly understood. Cells deficient in DNA methylation struggle to complete S phase suggesting a role for DNA methylation in regulating replication (Haruta et al., 2016; Jackson-Grusby et al., 2001; Jacob et al., 2015; Liao et al., 2015).

Gains and losses of the long arms of human chromosomes (Chr) 1, 9 and 16 are strongly correlated with hypomethylation and are overrepresented across many types of cancers as well as in aging cells (2012; Neve et al., 2006; Qu et al., 1999a; Qu et al., 1999b; Suzuki et al., 2002; Tsuda et al., 2002). Chromosomal rearrangements (whole arm deletions, isochores, multiradial structures and decondensations) centering on the pericentromeres of Chr 1, 9 and 16 have been reported in stimulated lymphocytes and fibroblasts from patients with the fatal genetic disease ICF syndrome (Ehrlich et al., 2001). ICF syndrome is caused by germline mutations in the de novo DNA methyltransferase, *DNMT3B*; the zinc finger protein, *ZBTB24*; the lymphoid specific helicase, *LSH/HELLS*; and *CDCA7*, which result in a striking depletion of methylation in the pericentromeres of these three chromosomes (de Greef et al., 2011; Ehrlich et al., 2001; Hernandez et al., 1997; Thijssen et al., 2015; Xu et al., 1999). The pericentromeres of these chromosomes are unique in that they contain long tracts of the closely-related repetitive sequences, Satellite 2 and 3 (SAT2/3)(Altemose et al., 2014; Cooke and Hindley, 1979; Jeanpierre et al., 1985; Moyzis et al., 1987).

SAT2/3 are enriched for tandem repeats of the pentamer, GGAAT, as well as diverged sequences, including <u>CG</u>GAT, and are densely methylated in healthy cells (Grady et al., 1992; Miller et al., 1974). Notably, hypomethylation of SAT2/3 is frequent across tumor types and is strongly correlated with a worse prognosis (Narayan et al., 1998; Ting et al., 2011). Structural studies have demonstrated that the G-rich sequence (GGAAT)_4_ folds into an unusually stable stem-loop structure (Catasti et al., 1994). Such non-canonical DNA structures are known to stall replication forks and lead to the formation of breaks and chromosomal rearrangements. Thus, we hypothesized that hypomethylation of SAT2 and SAT3 sequences leads to chromosomal abnormalities through the induction of replication stress (Leon-Ortiz et al., 2014).

Here we report that the human pericentromeric repeat sequence SAT2 poses challenges to the DNA replication machinery when hypomethylated. We have used well-characterized ICF patient and control cell lines to characterize the DNA damage response elicited at hypomethylated SAT2/3 sequences. We show that loss of methylation at SAT2, but not SAT3, is associated with increased frequencies of chromosomal abnormalities and DNA damage. We find that hypomethylation of SAT2/3 is associated with elevated levels of replication stress signaling and that chromosomal abnormalities involving SAT2 are enhanced by low levels of aphidicolin-induced replication stress. To investigate the basis for these chromosomal abnormalities, we developed a single-molecule approach employing DNA combing to directly visualize the impact of hypomethylation on replicating SAT2 sequences. Our analysis of replicating SAT2 molecules provides in vivo evidence that hypomethylation of SAT2 strongly decreases the efficiency of replicating these sequences, suggesting that hypomethylation results in the formation of barriers to the replication machinery. Consistent with the model we have proposed, we find increased levels of single-stranded DNA (ssDNA) binding protein RPA2 as well as asymmetric progression of sister replication forks within hypomethylated SAT2 sequences compared with normally methylated SAT2 sequences. Our data supports a model where chromosomal rearrangements at hypomethylated SAT2 sequences are triggered by replication stress and also provides potential mechanistic insight into the underlying causes of genomic instability in hypomethylated cells in diverse pathological conditions.

## Results

### Hypomethylation of SAT2, but not SAT3, is associated with chromosomal abnormalities and DNA damage

In the following experiments, we compare the well-studied lymphoblastoid cell line (LCL) derived from an ICF patient (DNMT3B^A603T/STP807ins^) (“ICF LCLs”) to control LCLs from the unaffected mother (DNMT3B^A603T/+^)(“Ctrl LCL”)(Ehrlich et al., 2001; Hansen et al., 1999). SAT2 and 3, which are present in multi-megabase (MB) long tracts in the pericentromeres of human chromosomes 1, 9 and 16 (Figure 1A), have been reported to be hypomethylated in ICF LCLs (Altemose et al., 2014; Ehrlich et al., 2008). We used methylation-sensitive southern blotting to confirm hypomethylation of SAT2 and 3 in ICF LCLs and normal methylation of these sequences in Ctrl LCLs (Supplemental Figure 1A). We used thin layer chromatography (TLC) to determine whether 5-methylcytosine levels were globally depleted in these cells. Genomic DNA was digested with the methylation-insensitive restriction enzymes MspI or Taq^*α*^I, which cleave DNA at the sequences C^CGG and T^CGA respectively. The resulting fragments, whose 5′ ends derive from the dinucleotide CpG and contain either C or 5mC, were end-labeled and digested to yield 5′ phosphorylated dNMPs that were resolved using TLC (Supplemental Figure 1B). Using this approach, we demonstrated that there is no difference in genome-wide methylation levels between Ctrl and ICF LCLs (Supplemental Figure 1B and C). Together, this demonstrates that the impact of *DNMT3B* mutations on methylation in ICF patients is restricted to limited portions of the genome, such as SAT2 and 3, while genome-wide methylation levels are not perturbed.

**Figure 1.**
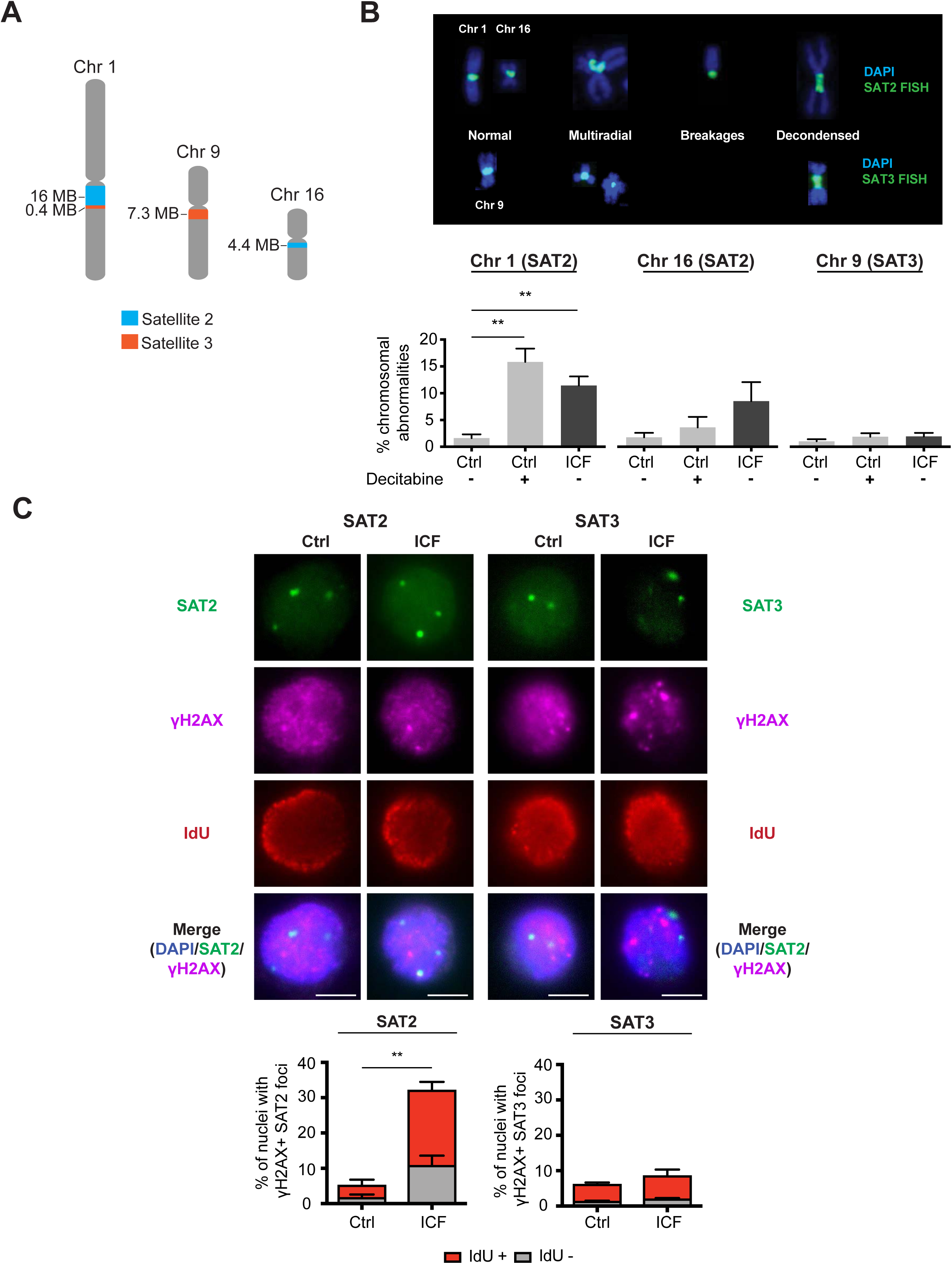
Hypomethylation of Satellite 2 (SAT2), but not Satellite 3 (SAT3), is associated with increased frequencies of chromosomal abnormalities and DNA damage. (A) The pericentromeres of Chr 1, 9, and 16 are composed of multi-megabase (MB) long tracts of densely methylated SAT2 and SAT3 sequences. (B) Hypomethylated SAT2 sequences in Immunodeficiency, <u>C</u>entromeric instability, and <u>F</u>acial anomalies (ICF) syndrome Lymphoblastoid cell lines (LCLs) show a higher frequency of chromosomal abnormalities (breakages, fusions, decondensation and multiradials) compared to normally methylated SAT2 sequences in control (Ctrl) LCLs. LCLs are treated with 2 µM Decitabine for 72 h to demethylate genome. (C) Hypomethylated SAT2 sequences in ICF LCLs show a higher frequency DNA damage (γH2AX) compared to normally methylated SAT2 sequences in Ctrl LCLs. SAT3 sequences do not exhibit a higher frequency of chromosomal abnormalities or DNA damage when hypomethylated. The majority of ICF LCLs with damaged SAT2 sequences are in S phase as determined by 5-chloro-2’-deoxyuridine (CldU) incoporation. Scale bar represents 5 µm. Mean values ±S.D. derived from at least three independent experiments. **P < 0.01; one-tailed Mann-Whitney U test.

As previously documented, we found a striking increase in chromosomal abnormalities (breakages, decondensations, fusions and multiradial chromosomes) involving hypomethylated SAT2 on Chr 1 (∼7-fold increase) and Chr 16 (∼5-fold increase) in ICF LCLs compared to Ctrl LCLs (Figure 1B).

Importantly, we found that genome-wide demethylation of Ctrl LCLs by treatment with the DNA methyltransferase inhibitor Decitabine for 72 h (2 µM Dec) induces similar levels of chromosomal abnormalities on Chr 1 and Chr 16. Surprisingly, we found that chromosomal abnormalities involving hypomethylated SAT3 sequences on Chr 9 occur much less frequently. Moreover, these abnormalities are only mildly elevated (<2-fold) in both ICF LCLs and Ctrl LCLs treated with Dec (Figure 1B). Although both SAT2 and SAT3 have frequently been described together as displaying hypomethylation-dependent instability (Jeanpierre et al., 1993), the decreased sensitivity of SAT3 to hypomethylation we observe is broadly consistent with the statistics presented in several papers (Ehrlich et al., 2006; Hernandez et al., 1997; Tuck-Muller et al., 2000).

Hypomethylation of somatic cells has been shown to result in elevated levels of DNA damage markers, such as *γ*H2AX (Haruta et al., 2016; Liao et al., 2015). We used immunofluorescence (IF) combined with fluorescence in situ hybridization (FISH)(Immuno-FISH) to ask whether DNA damage can be detected specifically at SAT2/3 sequences in hypomethylated cells. We find a six-fold enrichment of co-localization of *γ*H2AX foci with SAT2 FISH signals in ICF LCLs compared with Ctrl LCLs (Figure 1C). In contrast, SAT3 does not co-localize with *γ*H2AX in ICF patient cells providing further evidence that DNA methylation is not as critical for SAT3 stability as it is for the stability of SAT2 (Figure 1C).

### Chromosomal abnormalities and DNA damage at hypomethylated SAT2 are associated with replication stress

In order to test whether the chromosomal abnormalities at hypomethylated SAT2 sequences can be enhanced by replication stress, we examined metaphases of ICF LCLs treated with a low concentration of Aphidicolin (0.2 µM Aph) for 17 h. Low doses of Aph partially inhibit DNA polymerase and selectively impair the progression of replication forks through physiological barriers, such as non-B form DNA structures (Lukas et al., 2011). We find Aph increases (∼2-fold) the frequency of chromosomal abnormalities involving SAT2 in ICF LCLs, but not in Ctrl LCLs (Figure 2A). This suggests that SAT2 hypomethylation results in the formation of barriers to replication that are not present when these sequences are normally methylated. Aph treatment does not enhance chromosomal abnormalities at either normally methylated or hypomethylated SAT3 sequences indicating an lack of replication stress at SAT3 regardless of its methylation state. Furthermore, consistent with a role for replication stress in generating DNA damage at hypomethylated SAT2, the ATR target CHK1 is phosphorylated in ICF patient LCLs. In contrast, phosphorylation of the ATM target CHK2 is not detected (Figure 2B). Finally, we asked whether progression through S phase is associated with the generation of DNA damage. Since LCLs cannot be synchronized using a double-thymidine block (Chau et al., 2008), we treated cells with the thymidine analog 5-iodo-2’deoxyuridine (IdU) for 30 min to identify replicating cells. We find that over two-thirds of ICF LCLs with *γ*H2AX^+^ SAT2 foci are in S phase (Figure 1C).

**Figure 2.**
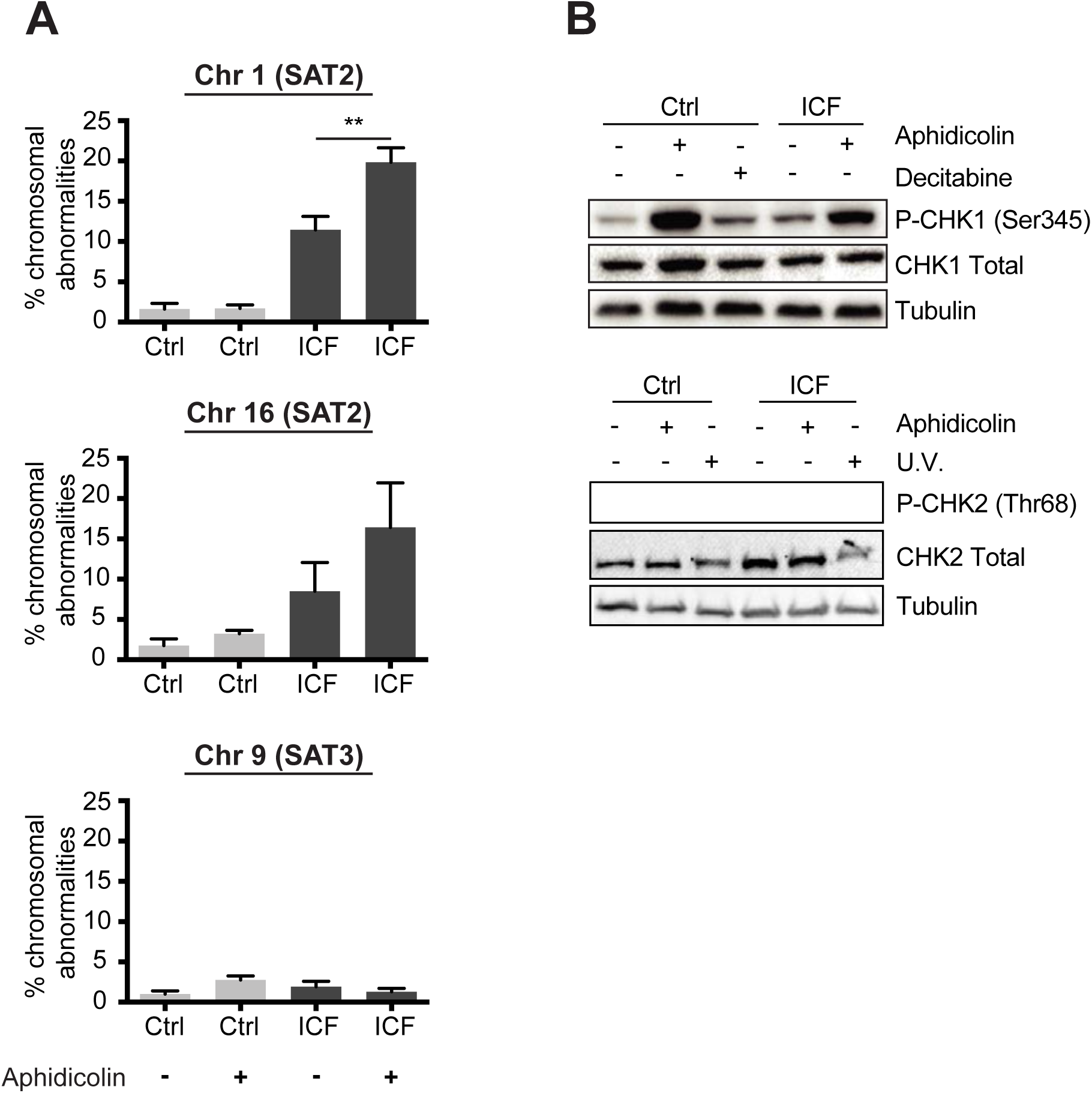
Chromosomal abnormalities and DNA damage at hypomethylated SAT2 are associated with replication stress. (A) Chromosomal abnormalities at hypomethylated SAT2 but not SAT3 are enhanced by treatment with low doses of the DNA polymerase inhibitor aphidicolin (0.2 μM Aph) for 17 h. (B) The ATR target CHK1 is phosphorylated (S345) in ICF patient LCLs, whereas phosphorylation of the ATM target CHK2 (T68) is not detected. Mean values ± SEM. derived from at least three independent experiments. **P < 0.01; one-tailed Mann-Whitney U test.

### Single-molecule analysis reveals that hypomethylation impairs replication of SAT2 sequences

The effect of hypomethylation on SAT2 replication has never been studied directly. Better tools are needed to study the replication profile of these sequences and to determine whether impaired fork progression and stalling underlie the increased frequency of chromosomal rearrangements in hypomethylated cells. To obtain a better understanding of how hypomethylation impacts replication, we combined DNA combing with SAT2 FISH to examine the progress of replication forks through SAT2 at the level of a single DNA molecule (Anglana et al., 2003; Bianco et al., 2012). We utilize this approach throughout the work described here to test the contribution of DNA methylation on replication fork progression.

Briefly, cultured cells are pulsed with the thymidine analog IdU, and DNA replicating during pulse periods becomes labeled with this nucleoside. Genomic DNA is extracted from labeled cells, embedded in agarose plugs, stretched on vinyl silane coated glass coverslips using Genomic Vision’s Molecular Combing System (MCS), and then denatured (Anglana et al., 2003; Bianco et al., 2012). Incorporation of IdU is visualized using a fluorescent anti-IdU antibody (red). SAT2-containing DNA molecules are identified using biotinylated FISH probes (blue) that are specific for the SAT2 arrays found on Chr 1 and 16 (Figure 3A-B) (Tagarro et al., 1994). Genome-wide rates of replication are calculated using non-denatured whole-genome fiber spreads prepared in parallel and stained with anti-IdU. We have found that DNA fibers stretched using the MCS do not require denaturation in order for anti-IdU antibodies to bind, allowing individual DNA molecules to be visualized simply by incubation with the DNA binding dye, YOYO-1 (green)(Figure 3B).

**Figure 3.**
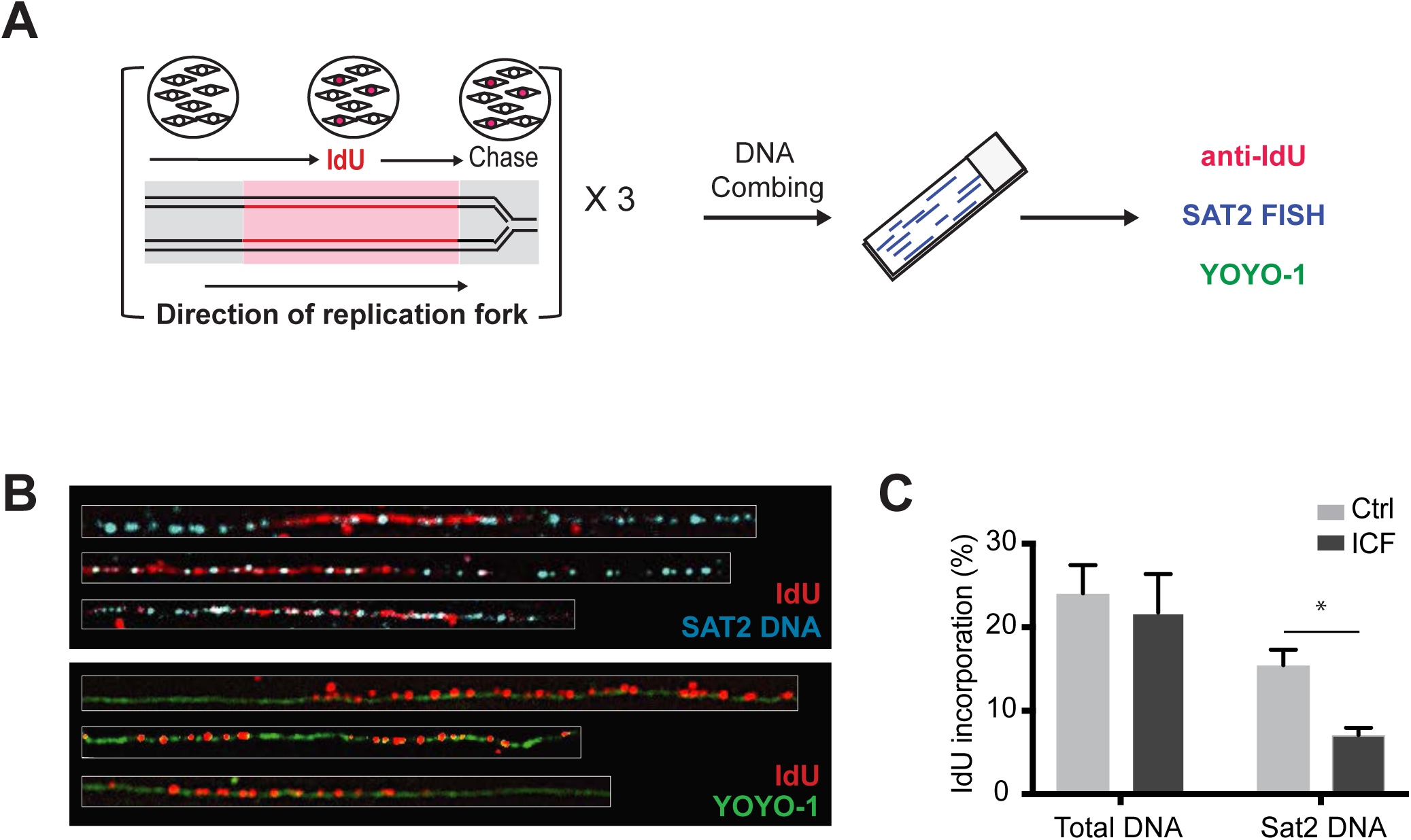
SAT2 hypomethylation is associated with decreased replication efficiency. (A) Schematic for using Replication Combing Assay (RCA) with SAT2 FISH to study the impact of hypomethylation on replicating SAT2 sequences: Cells are pulsed with 5-iodo-2’-deoxyuridine (IdU)-containing media followed by a 3 h chase with normal media. Purified genomic DNA is stretched on vinyl silane coated glass coverslips and stained with an antibody specific for IdU (red). SAT2 is identified using biotinylated fluorescence in situ hybridization (FISH) probes (blue), and YOYO-1 is used to identify DNA fibers (green). (B) Examples of stained and stretched i) SAT2 DNA molecules labeled with IdU and SAT2 FISH probes, and ii) DNA molecules labeled with IdU and YOYO-1. (C) Hypomethylation of SAT2 sequences is associated with reduced replication efficiency, while genome-wide replication efficiency is unaffected. Total DNA replication efficiency = (Σ [length IdU] / Σ [length DNA]). SAT2 replication efficiency *=* (Σ [length IdU] / Σ [length SAT2]). Mean values ± SEM. derived from at least three independent experiments. *P ≤ 0.05; one-tailed Mann-Whitney U test.

Under the conditions used, 24% of total DNA becomes labeled with IdU (*Σ [length IdU] /Σ [length total DNA]*) in Ctrl LCLs, whereas only 16% of SAT2 sequences in Ctrl LCLs become labeled with IdU (*Σ [length total IdU] /Σ [length total SAT2]*) (Figure 3C). To determine whether hypomethylation affects the efficiency of SAT2 replication, we have used this approach to determine the efficiency of IdU incorporation in ICF LCLs. In support of our hypothesis, we find SAT2 hypomethylation results in ∼2-fold lower incorporation of IdU at SAT2 (decreases from 16% to 8%), indicating that hypomethylation strongly diminishes the efficiency of replicating SAT2 (Figure 3C). This is not due to a general effect on DNA replication, as genome-wide rates of nucleoside incorporation are not altered in these cells (Figure 3C).

### Increased levels of single-stranded DNA (ssDNA) binding protein RPA2 accumulate at hypomethylated SAT2 sequences

While diminished replication efficiency is consistent with elevated rates of fork stalling at hypomethylated SAT2 sequences, it could also be caused by slower movement of the replication fork through these sequences as well as by decreased origin firing. Therefore, we are taking several approaches to definitively address whether fork stalling occurs more frequently at hypomethylated SAT2 sequences. Replication fork stalling leads to the functional uncoupling of the replicative helicase from the stalled DNA polymerase (Byun et al., 2005). This leads to the generation of single-stranded DNA (ssDNA) at the fork junction and the accumulation of focal concentrations of the ssDNA binding proteins, RPA1/2/3 (Balajee and Geard, 2004; Ouyang et al., 2013). We use Immuno-FISH to ask whether RPA2 foci are elevated at SAT2 in ICF LCLs compared to Ctrl LCLs. We find that ICF LCLs exhibit ∼2.5-fold increase in the percentage of nuclei containing SAT2 foci that co-localize with RPA2 compared to Ctrl LCLs (Figure 4A-B). This data indicates that more ssDNA is present at hypomethylated SAT2 sequences in ICF LCLs compared with Ctrl LCLs, which is consistent with increased frequency of fork stalling at these sequences.

**Figure 4.**
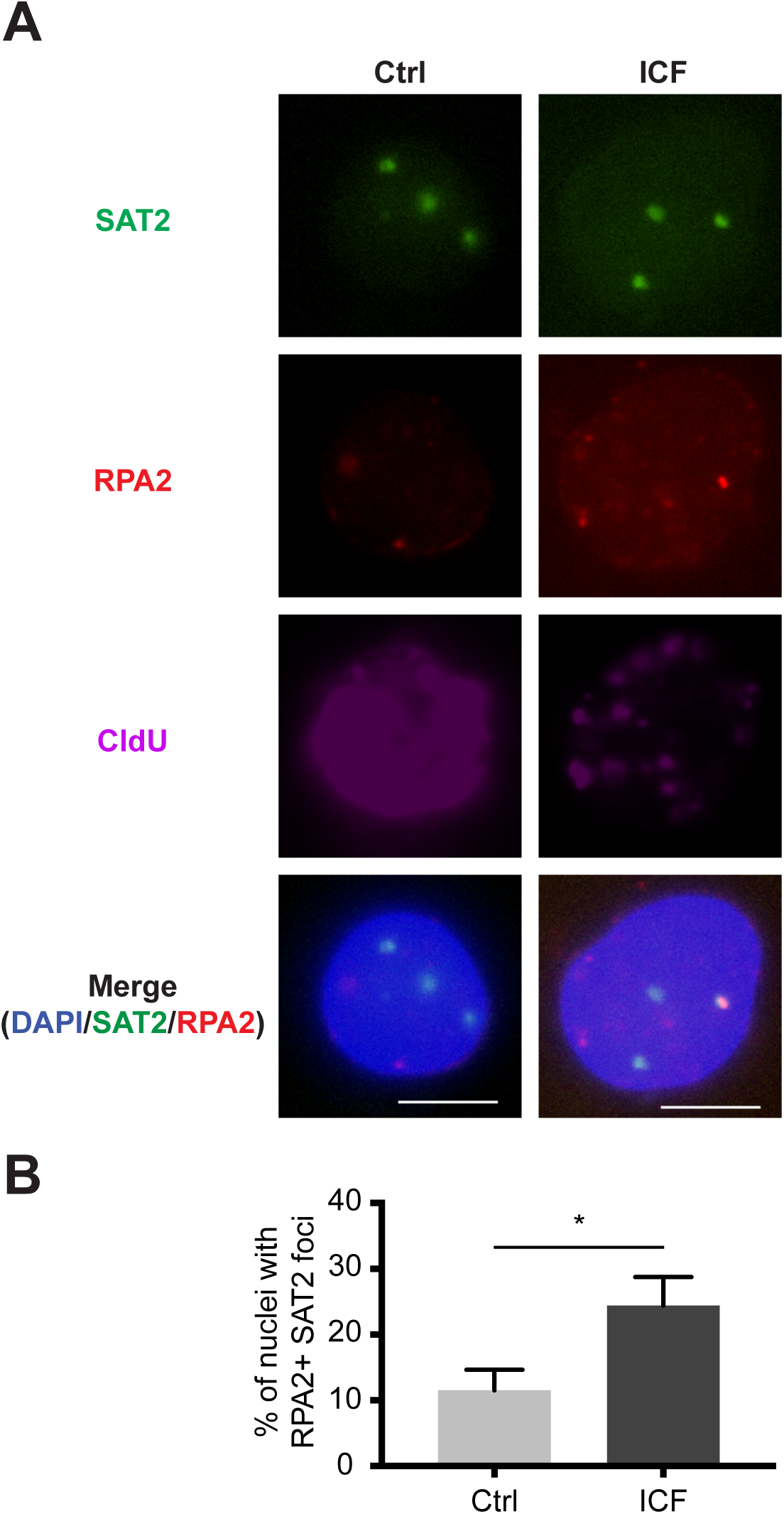
RPA2 accumulates at hypomethylated SAT2 in replicating cells. (A) Immunofluorescence (IF)-FISH images showing colocalization of RPA2 foci with SAT2 foci in ICF LCLs but not Ctrl LCLs. LCLs were pulsed with 25 μM 5-chloro-2’-deoxyuridine (CldU) for 30 min to identify cells in S phase. CldU (purple) and RPA2 (red) were detected with fluorescent antibodies, SAT2 (green) was detected with a fluorescent FISH probe, and DNA (blue) was counterstained with DAPI. Scale bars, 5 μm. (B) Quantification of IF-FISH data presented in (A). Results are expressed as the percentage of cells presenting at least one SAT2 foci overlapping with an RPA2 foci, and represent the mean of three biological replicates. A minimum of 40 cells were analyzed per replicate. Mean values ±S.D. derived from at least three independent experiments. *P ≤ 0.05; one-tailed Mann-Whitney U test.

### Hypomethylation of SAT2 results in increased frequency of fork stalling

In order to definitively address whether SAT2 hypomethylation increases the frequency of replication fork stalling, we have adapted our single-molecule approach to allow us to assess whether bidirectional replication forks move at equal rates in both leftward and rightward directions. In this modified procedure, cultured cells are pulsed sequentially with thymidine analogs, 5-chloro-2’-deoxyuridine (CIdU) and then ldU. Genomic DNA is purified, stretched on silanized glass coverslips and denatured as described. Stretched DNA molecules are stained with antibodies that distinguish CldU (red) from IdU (green). SAT2-containing DNA molecules are identified by biotinylated FISH probes (blue) (Figure 5A) (Tagarro et al., 1994). A key challenge is the identification of conditions that allow sufficient numbers of dual-labeled SAT2 molecules to be obtained for analysis. The optimized procedure uses labeling periods of 15 min with CldU and followed by 15 min with IdU. SAT2 is an abundant repeat, but represents only 1-2% of the human genome (Miga, 2015). To facilitate the identification of dual-labeled SAT2 molecules in whole genome spreads, we use digital scanning and image analysis software capable of capturing molecules on each coverslip and recognizing and classifying signals of interest (FiberVision Automated Scanner and FiberStudio Analysis Software, Genomic Vision). Evidence for fork stalling in SAT2 will be provided by determining the frequency of molecules that have initiated replication within a stretch of SAT2 and show a nonterminal CldU (red) segment flanked by asymmetric incorporation of ldU (green) on either side (Figure 5B). This will indicate that the fork progressed on one side during the ldU pulse, while it stalled on the other side. An increase in the frequency of asymmetric forks at hypomethylated SAT2 sequences in ICF LCLs compared with the frequency in control LCLs would indicate that hypomethylation of SAT2 sequences results in fork stalling.

**Figure 5.**
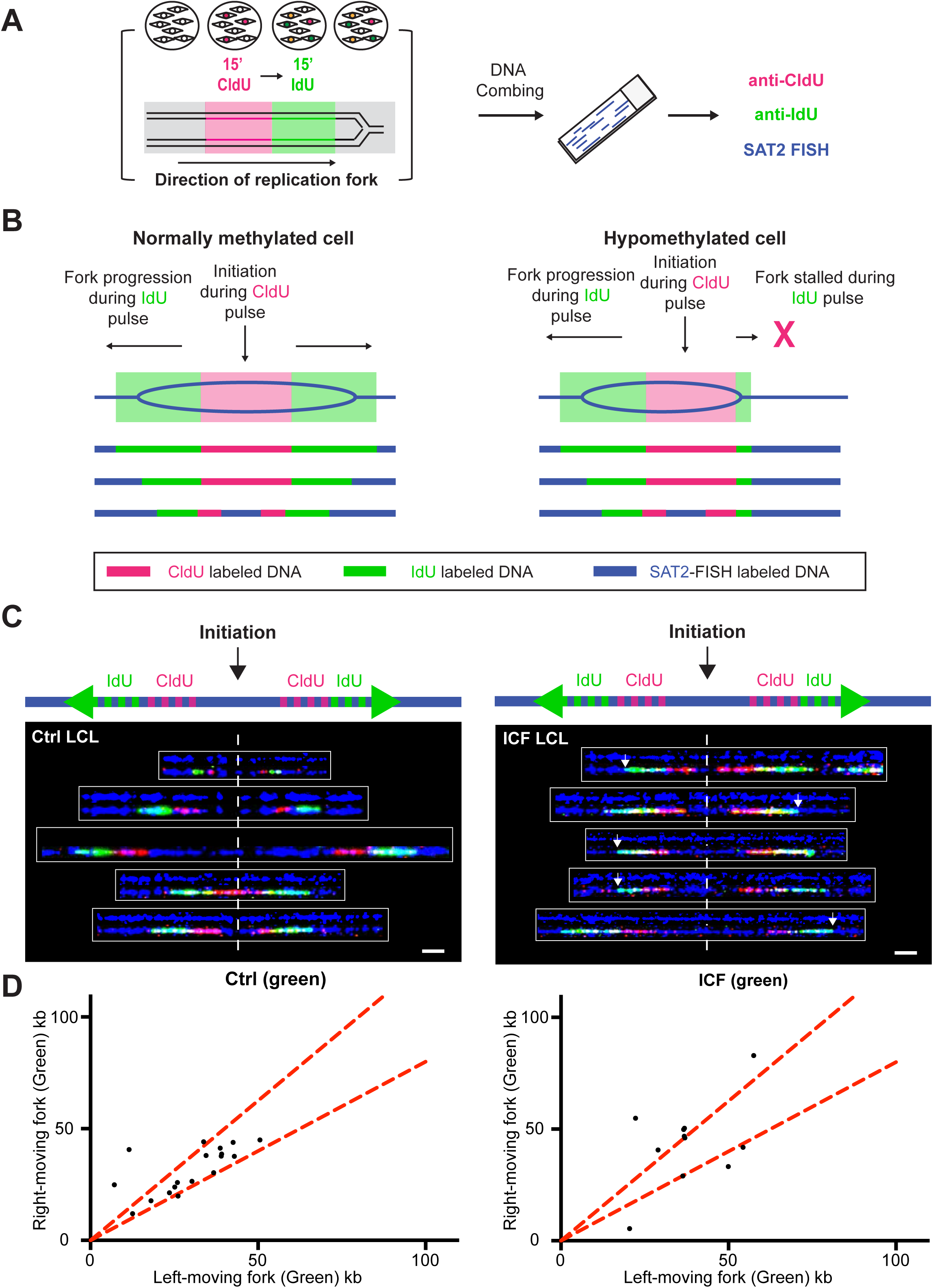
Single molecule analysis demonstrates increased replication fork stalling in hypomethylated SAT2 molecules. (A) Schematic for DNA combing with SAT2 FISH to study the impact of hypomethylation on replicating SAT2 sequences. Cells are pulsed sequentially with CldU and then IdU-containing media. Purified genomic DNA is stretched on silanized glass coverslips and then stained with antibodies specific for CldU (red) and IdU (green). SAT2 is identified using biotinylated FISH probes (blue). (B) The predicted pattern of labeling by RCA in normally methylated and hypomethylated SAT2 sequences is shown below an illustration of bidirectional fork progression. (C) Representative images of sister replication forks in Ctrl and ICF LCLs. Scale bar represents 10 µm. White arrows denote unstable and/or stalled replication forks. (D) Evidence for fork stalling in molecules that initiated replication within SAT2 sequences. Quantitation of asymmetry between left-moving and right-moving sister forks demonstrates that 91% of bidirectional replication forks found in Sat2 are asymmetric in ICF LCLs compared with 22% in Ctrl LCLs. The central area between the red dotted lines contain sister forks with less than 20% length difference.

Using this approach, we are able to identify dual-labeled molecules that have initiated replication within stretches of SAT2 indicating that SAT2 is replicated from bidirectional origins contained within the repeat array (Figure 5C). Of these bidirectional replication forks, 91% were highly asymmetrical in ICF LCLs, compared to only 22% in Ctrl LCLs (Figure 5D) demonstrating that the loss of methylation in ICF patient cells results in increased replication fork stalling and/or replication collapse at SAT2 sequences

## Discussion

Here we show that hypomethylated SAT2 sequences present challenges to the DNA replication machinery. We show that loss of methylation at SAT2, but not SAT3, is associated with increased frequencies of chromosomal abnormalities and DNA damage (Figure 1). We also show that hypomethylation of SAT2/3 is associated with replication stress signaling and that chromosomal abnormalities involving SAT2 are enhanced by low levels of aphidicolin-induced replication stress (Figure 2). Our single-molecule approach revealed that hypomethylation of SAT2 impairs the efficiency of replicating SAT2 (Figure 3). Finally, consistent with our model of hypomethylation driven fork stalling, we find increased levels of single-stranded DNA (ssDNA) binding protein RPA2 as well as asymmetric progression of sister replication forks within hypomethylated SAT2 sequences compared with normally methylated SAT2 sequences (Figures 4-5). Our data supports a model where chromosomal rearrangements at hypomethylated SAT2 sequences are triggered by replication stress.

Several interesting questions still remain to be answered. It is not clear how DNA methylation promotes efficient replication of the genome. One possibility is that dynamic changes in DNA methylation directly regulate isomerizations between classic helical DNA (B-form DNA) and non-canonical DNA structures. Another possibility is that DNA methylation represses transcription of SAT2 and prevents the formation of RNA:DNA hybrids, another type of non-canonical DNA structure known to stall replication forks (Crossley et al., 2019; Gan et al., 2011; Hamperl et al., 2017). Recent reports from several groups have demonstrated that non-coding RNAs are transcribed from pericentromeric satellite sequences in G2/M and associate in cis to form RNA:DNA hybrids (Johnson et al., 2017; Velazquez Camacho et al., 2017). In light of these new findings, it will be necessary to investigate whether hypomethylation selectively derepresses transcription of SAT2, and not SAT3. It should be noted that this possibility is not mutually exclusive with the proposal that DNA methylation directly regulates the formation of non-canonical DNA structures as formation of an RNA:DNA hybrid may allow non-B DNA structures to fold on the strand of DNA that is not part of the hybrid.

It is also not clear why SAT2 displays strong hypomethylation-induced genomic instability compared to SAT3 when both are enriched for sequences that have the potential to fold into non-canonical DNA structures (Catasti et al., 1994; Grady et al., 1992). One possible explanation is that SAT2 has a higher abundance of sequences that can fold into non-canonical DNA structures. A second possibility is that SAT2 contains sequences and structures that are regulated by DNA methylation that are not present in SAT3. Further studies to identify sequences with secondary structure forming potential in SAT2 and SAT3 and to test the impact of methylation on the formation of non-canonical structures should provide answers to these questions.

In support of the idea that methylation has a distinct function at SAT2, we find that the methylatable dinucleotide, CpG, is unusually abundant at SAT2. Due to the inherent mutability of methylated cytosine, the abundance of the dinucleotide CpG in the mammalian genome is much lower than the 4% predicted based on GC content (Lander et al., 2001). Deamination of cytosine results in the production of uracil, which is recognized as abberant and efficiently repaired back to cytosine. However, deamination of methyl cytosine results in the production of thymine. This leads to transition mutations (C→T), the loss of methylated CpGs, and ultimately yields the observed genome-wide levels of CpG abundance of ∼1% (Bird, 2002). This scarcity of CpGs in the genome is not uniform and kilobase (kb) long stretches of CpG rich sequences (CpG islands (CGIs)) persist, ostensibly reflecting the fact that these regions are not methylated in the germline (Bird, 2002). We analyzed the CpG content in SAT2 and SAT3 using whole genome shotgun reads previously classified as SAT2 or SAT3 in the HuRef Genome (Altemose et al., 2014). Our analysis shows that the frequency of CpGs is significantly higher in SAT2 (∼6.2%) than the 4% expected based on GC content. This value is especially striking considering that the vast majority of these CpGs are methylated and thus are subject to attrition. In contrast to SAT2, SAT3 contains 3% CpGs. This is higher than genome-wide levels of CpG abundance, but not as high as the levels seen in SAT2. Taken together this suggests that SAT2 has evolved a unique role for regulation by methylation that may not be present at SAT3.

The findings we present here indicate that impaired replication underlies the formation of DNA damage and chromosomal aberrations seen at hypomethylated SAT2 sequences. This work suggests a mechanistic basis for how the loss of DNA methylation may contribute to genomic instability in diverse pathological conditions. Future studies will be aimed at determining whether the hypomethylation frequently documented in aging and cancer cells is a critical factor driving the replication stress that is believed to promote both aging and tumor evolution (Macheret and Halazonetis, 2015; Zeman and Cimprich, 2014).

## Supplemental Figure Legends

**Supplemental Figure 1.**
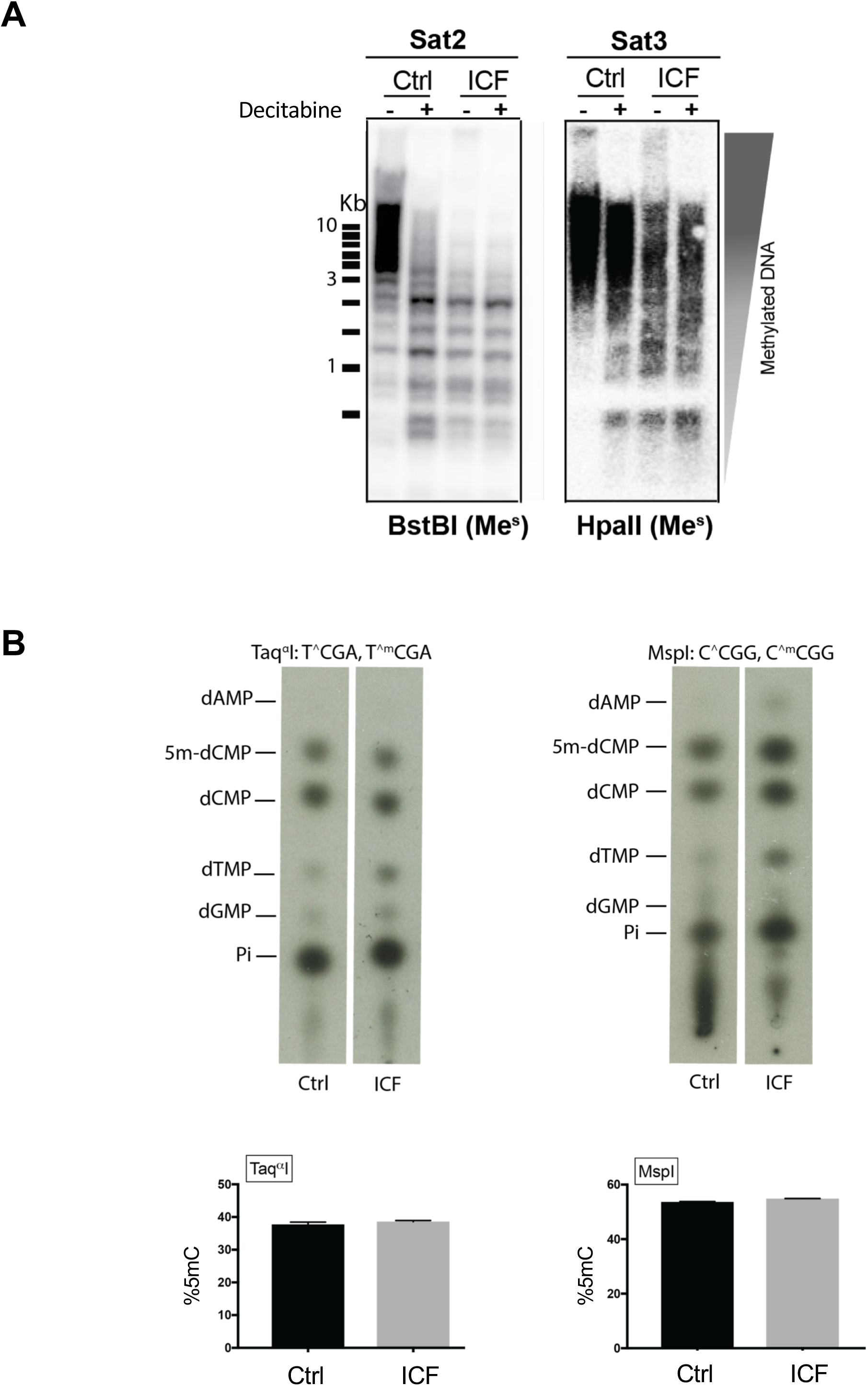
ICF LCLs are hypomethylated at SAT2 and SAT3 compared with Ctrl LCLs, but genome-wide levels of cytosine methylation do not differ between these cell lines. (A) Digestion of genomic DNA with methylation-sensitive enzymes (BstBI for SAT2, HpaII for SAT3) shows DNMT3B^A603T/STP807ins^ ICF LCLs are hypomethylated at SAT2 and SAT3 compared with control (Ctrl) LCLs from unaffected mother DNMT3B^A603T/+^. (B) *Upper panels* Genome-wide levels of cytosine methylation do not differ between Ctrl and ICF LCLs. Genomic DNA was purified from LCLs and cleaved with methylation insensitive restriction enzymes MspI and Taq^*α*^I. The fragments were end-labeled, digested to 5′ dNMPs, and resolved using thin layer chromatography (TLC). *Lower panels* Quantification of the relative abundance of dCMP and 5m-dCMP. Mean values ±S.D. derived from at least three independent experiments.

## Methods

### Southern Blot

500 ng of genomic DNA was digested with the methylation sensitive enzymes, BstBI (NEB R0519L) or HpaII (NEB R0171S), and run on a 0.8% agarose gel. Gel was depurinated in 0.24 M HCl, denatured in 1.5 M NaCl / 0.5 M NaOH, and neutralized in 1.5 M NaCl / 1M Tris, pH 7.0. After equilibration in 10X SSC buffer, DNA was transferred overnight onto a nylon membrane and crosslinked with UV (1200 J/cm^2^). SAT2 and SAT3 DNA was detected with end-labeled oligonucleotides (SAT2: TCGAGTCCATTCGATGAT or SAT3: TCCACTCGGGTTGATT) (Tagarro et al., 1994). Membranes were exposed for at least 12 h using a Biomax MS Film at - 80 °C.

### Analysis of 5mC levels using thin-layer chromatography

Nuclei were prepared by resuspending cells in 1 mL NPB [320 mM sucrose, 10 mM Tris, pH 7.5, mM MgCl_2_, 1% Triton X-100, with 4 µg RNAseA/ml (Qiagen 158922)] and placing cells on ice for 20 min. Cells were spun at 1300 g for 15 min at 4 °C and then washed once in NPB. Nuclei were lysed in 650 µl of 1X LB (10 mM Tris, pH 8.0, 300 mM sodium acetate, pH 7.2, 0.5% SDS, 20 mM EDTA, 8 ug RNAseA/ml) and placed at 37 °C for 1 h. 50 µg Proteinase K (Roche 3115801001) was added and incubated overnight at 55 °C. An extra 50 µg/ml Proteinase K was added in the morning and the samples were left at 55 °C for 5 h. Samples were extracted with equal volumes of phenol, phenol:chloroform:isoamyl alcohol (25:24:1), and chloroform:isoamyl alcohol (24:1) and then precipitated with 2 volumes of ethanol. Genomic DNA was washed twice with 1 mL of 72% EtOH, dried, and resuspended in 10 mM Tris, 0.1 mM EDTA, pH 8.0 and allowed to resuspend overnight at 32 °C.

2 µg of genomic DNA was digested with 100 units of MspI (NEB R0106M) or Taq^*α*^I (NEB R0149M) and 4 µg of RNaseA (Qiagen 28306) overnight as per the manufacturer’s instructions. An extra 100 units of restriction enzyme was added in the morning and incubations were continued for 6 hours. 10 units of calf intestinal phosphatase (CIP) (NEB M0290L) was added and incubated for 1 h at 37 °C. DNA was purified using Qiaquick Nucleotide Removal Kit (Qiagen) as per the manufacturer’s instructions. 400 ng of eluted DNA fragments were end-labeled with T4 Polynucleotide Kinase (T4 PNK) (NEB M0201L) and 10 µCi of [*α*-^32^P]-ATP (Perkins Elmer NEG002A) for 1 h at 37 °C. Labeled fragments were precipitated by the addition of 30 µg of linear polyacrylamide, 1/10 volume of 3 M sodium acetate, pH 7.2 and 2.5 volumes of 100% ethanol at left at −80 °C for 1 h. Samples were spun at 14,000 rpm for 20 minutes at 4 °C and washed twice with 70% ethanol at 25 °C. Pellets were resuspended in 30 mM Tris, pH 8.9, 15 mM MgCl_2_, 2 mM CaCl, with 10 µg of DNaseI (Worthington, LS00631) and 10 µg SVPD (Worthington, LS003926) and incubated for 3 h at 37 °C. 2 µl were spotted on cellulose TLC plates (20 cm x 20 cm, VWR EM1.05718.0001) along with 10 µg of dNMP standards and developed in isobutyric acid:H_2_O:NH_3_ (66:20:1). Plates were analyzed by phosphorimager scanning using ImageQuant TL software (GE Healthcare Life Sciences). The low-level labeling of other nucleotides reflects DNA shearing or contaminating endonucleolytic activity.

### Metaphase fluorescence *in situ* hybridization (FISH)

LCLs were harvested while in exponential phase, incubated with Demecolcine solution (Sigma D1925) for 30 min at room temperature (RT), swelled in 75 mM KCl for 30 min at RT, fixed in ice cold methanol and acetic acid solution (3:1 ratio) and dropped onto a glass slide. Slides were dehydrated in 70%, 90%, and 100% ethanol baths, and chromosomes were denatured for 2 min in 70% formamide / 2X SSC at 80 °C. Slides were immediately transferred to a series of ice-cold, 70%, 90%, and 100% ethanol baths. Hybridization was performed overnight at 37 °C in a humid chamber.

The hybridization solution contains 1 µg/ml FISH probe (SAT2: Biotin-TCGAGTCCATTCGATGAT-Biotin or SAT3: Biotin-TCCACTCGGGTTGATT-Biotin), 50 µg/mL denatured salmon sperm DNA (ThermoFisher AM9680) in 25% formamide, 4X SSC, 1 mM EDTA in 1X PBS.

After hybridization, the slides were washed in 2X SSC and biotin was detected with Streptavidin AF 488 (ThermoFisher S-32354; Lot# 1571714, 1977355). Slides were mounted and counterstained in Vector shield mounting medium with DAPI (Vector laboratories, H-1200). A minimum of 100 metaphases were analyzed using a Metafer slide scanning platform (Metasystem).

### Immunofluorescence (IF) combined with DNA FISH (Immuno-FISH)

Exponentially growing lymphoblastoid cells lines (LCLs) were seeded in wells containing poly-L-lysine (Sigma P6282) coated coverslips and incubated with 25 µM 5-chloro-2’-deoxyuridine (CldU) (MP Biomedicals 0210547891) (for RPA2 detection) or 25 µM 5-iodo-2’-deoxyuridine (IdU) (MP Biomedicals 0210035701) (for γH2AX detection) for 30 min at 37 °C. Cells were permeabilized in ice-cold CSK Buffer (10 mM HEPES-KOH, pH 7.4, 300 mM sucrose, 100 mM NaCl, 3 mM MgCl_2_) containing 0.5% Triton X-100 for 5 min, washed in CSK Buffer for 5 min, and fixed with 2% paraformaldehyde (PFA) / 2% sucrose for 10 min, all on ice. Following one quick rinse with 1X PBS, cells were permeabilized in cold 0.4% Triton X-100 / 1X PBS for 10 min on ice and rinsed once with 1X PBS. Cells were blocked for 30 min at RT with Blocking Buffer (2.5% BSA, 0.1% Tween, 10% FBS in 1X PBS). They were then incubated with either mouse-anti-RPA2 antibody (Abcam ab2175; Lot# GR323938-4) or rabbit anti-γH2AX (Cell Signaling 9718) diluted 1:200 in Blocking Buffer for 1 h at RT. After three washes in Wash Buffer (0.25% BSA, 0.01% Tween, 1% FBS in 1X PBS) for 5 min, cells were incubated with either goat anti-mouse AF 546 (ThermoFisher A11030; Lot# 1736960 and 1904466) or goat anti-rabbit AF 647 (ThermoFisher A21245) diluted 1:1000 in Blocking Buffer for 1 h at RT protected from light (for this and all subsequent steps). They were then washed three times in Wash Buffer for 5 min. Cells were post-fixed with 2% PFA / 2% Sucrose for 10 min on ice, washed once in 1X PBS, and incubated with 4 mg/mL RNase A (Qiagen 158922) diluted 1:286 in 2X SSC for 30 min at 37 °C. Cells were re-permeabilized with cold 0.7% Triton X-100 / 0.1M HCl for 10 min on ice, rinsed once with 1X PBS, and denatured in 2M HCl for 30 minutes at RT. After three quick rinses in cold 1X PBS, cells were incubated with FISH Hybridization Buffer [10 ng/uL FISH probe (Sat2: Biotin-TCGAGTCCATTCGATGAT-Biotin, Sat3: Biotin-TCCACTCGGGTTGATT-Biotin), 100 ng/uL denatured salmon sperm DNA (ThermoFisher AM9680), 5X Denhart’s solution, 10% dextran sulfate, 5X SSC, and 50% formamide] for 12-16 h in a dark and humid chamber.

Following hybridization, coverslips were washed three times in 2X SSC + 0.1% Tween for 5 min at RT and blocked with Blocking Buffer for 30 min at RT. Subsequently, cells were incubated with Streptavidin AF 488 (ThermoFisher S-32354; Lot# 1571714, 1977355) diluted 1:1000 in Blocking Buffer for detection of biotinylated FISH probes, and CldU primary antibody (rat anti-BrdU, Biorad OBT0030S; Lot# 0109) or IdU primary antibody (mouse anti-BrdU, BD Biosciences 347580; Lot # 7324574, 6334742) diluted 1:20 in Blocking Buffer. Following three washes with Wash Buffer for 5 min, cells were incubated with CldU secondary antibody (goat anti-rat AF647, ThermoFisher A-21247; Lot# 1696458, 200538) or IdU secondary antibody (goat anti-mouse AF 546, ThermoFisher A-11030; Lot# 1736960, 1904466) diluted 1:1000 in Blocking Buffer, and washed three times with Wash Buffer for 5 min. Coverslips were washed once in 1X PBS, once in distilled water, and mounted with DAPI diluted 1:100 in Prolong Gold (ThermoFisher P36930). Images were captured using a Nikon Eclipse T_i_ inverted microscope and results are expressed as percentage of cells presenting at least one SAT2 or SAT3 focus overlapping with γH2AX or RPA2 foci.

### Western Blot

Cells were lysed in 150 mM NaCl, 50 mM Tris pH 7.4, 1% NP40 supplemented with 1X protease inhibitor (Roche) and 1 mM sodium fluoride, 1 mM sodium orthovanadate, 25 mM B-glycerophosphate for 30 min on ice, sonicated 5 times for 30 s, and spun at 10,000 RCF for 10 min. Protein concentration was determined by Bradford assay. Following denaturation, 50 µg of protein was separated on a 10% acrylamide gel and transferred onto a nitrocellulose membrane. For the detection of phospho-CHK1 and phospho-CHK2, membranes were blocked in 5% BSA, 1X TBS, 0.1% (v/v) Tween for 1 h and detection was carried out overnight at 4 °C with rabbit anti-phospho-Chk1 (Ser345) mAb (133D3) (Cell Signaling 2348; Lot 15) or rabbit anti-Phospho-Chk2 (Thr68) (C13C1) Rabbit mAb (Cell Signaling 2197). Membranes were stripped and blocked in 5% milk in 1X TBS with 0.1% (v/v) Tween and detection of total CHK1 or CHK2 proteins was carried out for 1 h at RT using mouse-anti-Chk1 mAb (2G1D5) (Cell Signaling 2360; Lot 3) or mouse anti-Chk2 mouse mAb (BD Biosciences 611570). As a loading control, Tubulin was detected using rabbit Anti-Tubulin Alpha Antibody, C-Terminal (Sigma SAB450087; Lot 310379). All antibodies were detected using either goat anti-mouse HRP (Sigma A3682; Lot 071M4782) or goat anti-rabbit HRP (Sigma A0545, Lot 111M4750) conjugated secondary antibodies. Imaging was done using a ChemiDoc apparatus (Biorad) following incubation in Clarity^TM^ Western ECL Substrate (Biorad 17005060).

### DNA Combing Combined with FISH and IF

#### CldU/IdU Labeling and DNA Combing

To assess replication efficiency, exponentially growing LCLs were pulsed with 25 µM IdU for 1 h, harvested by centrifugation at RT, washed once in RT 1X PBS, and then incubated with pre-equilibrated media without IdU for 3 h (chase). This pulse-chase was repeated a total of three times.

To assess fork stalling, exponentially growing LCLs were sequentially labeled by incubation with 25 µM CldU (15 min) and then 25 µM IdU (15 min).

Cells were harvested by centrifugation, washed once in ice cold 1X PBS, and embedded in 1% Seaplaque low melt agarose (Lonza 50110) plugs at a concentration of 0.5×10^6^ cells/plug. Genomic DNA was extracted by incubating plugs overnight in Lysis Buffer [0.5M EDTA, pH 8.0, 1% N-lauroylsarcosine sodium with 1 mg/ml Proteinase K (Sigma 3115801001)]. Plugs were washed three times in Wash Buffer (10 mM Tris, pH 8, 1 mM EDTA) for 1 h and one time for 3 h at 4 °C, protected from light. Subsequently, plugs were equilibrated in 0.5 M MES, pH 5.5 (Calbiochem 475893) for 15-30 min, and melted at 70 °C for 30 min. DNA was equilibrated at 42 °C for 5-10 min, incubated with 1.5 - 3 µl GELase equilibrated in 50 µl of MES (Fisher G09100) overnight at 42 °C, and combed onto CombiCoverslips (Genomic Vision COV-001) using a FiberComb apparatus (Genomic Vision MCS-001) at RT. Following combing, coverslips were desiccated for 2 h at 65 °C.

#### FISH and Detection by IF

Coverslips were dehydrated in 70% ethanol, 95% ethanol, and 100% ethanol for 3-5 min each. Coverslips were then denatured in Denaturation Buffer (0.1 M NaOH, 70% ethanol) for 10 min and fixed in Fixation Buffer (0.1 M NaOH, 70% Ethanol, 2% Glutaraldehyde) for 5 min at RT. Coverslips were dehydrated in 70% ethanol, 95% ethanol, and 100% ethanol for 3 min each and then air dried for 5 min. They were then incubated with FISH Hybridization Buffer [1 M NaCl, 50% formamide, 1% SDS, 5 mM Tris, pH 7.4) containing 100 µg/mL denatured salmon sperm DNA and 1 µg/mL SAT2 FISH probe (Biotin-TCGAGTCCATTCGATGAT-Biotin) for 12-16 h in a dark and humid chamber.

Following hybridization, coverslips were washed 3 times in 2X SSC with 0.1% Tween for 5 min and then rinsed once in 1X PBS. Coverslips were then blocked with BlockAid Blocking Solution (ThermoFisher B10710) for 20 min at RT. Signals were developed by sequential incubations in the dark with the following antibodies diluted in BlockAid, each followed by 3 washes in 1X PBS, 0.1% Tween for 5 min:

##### To assess replication efficiency of SAT2

(1) Streptavidin AF 488 (ThermoFisher S-32354; Lot# 1571714, 1977355) diluted 1:100 for 20 min, (2) biotinylated anti-streptavidin (Vector BA-0500; Lot y1002) diluted 1:50 for 20 min, (3) Streptavidin AF 488 diluted 1:100 for 20 min, (4) biotinylated anti-streptavidin diluted 1:50 and IdU primary antibody (mouse anti-BrdU, BD Biosciences 347580; Lot # 7324574, 6334742) diluted 1:25 for 1 h, (5) Streptavidin AF 488 and IdU secondary antibody (goat anti-mouse AF 546, ThermoFisher A-11030; Lot# 1736960, 1904466) diluted 1:50 for 30 min - 1 h. Coverslips were briefly rinsed in ddH_2_0 and mounted with Prolong Gold.

##### To assess replication efficiency of total DNA

(1) IdU primary antibody (mouse anti-BrdU, BD Biosciences 347580; Lot # 7324574, 6334742) diluted 1:25 for 1 h, (2) IdU secondary antibody (goat anti-mouse AF 546, ThermoFisher A-11030; Lot# 1736960, 1904466) diluted 1:50 for 30 min - 1 h. Coverslips were briefly rinsed in ddH_2_0 and mounted with Prolong Gold supplemented with 0.1% (v/v) YOYO-1 Iodide (ThermoFisher Y3601; Lot #1596076).

To estimate SAT2 and total DNA replication efficiency, images were captured using a Nikon Eclipse T_i_ inverted microscope. Tract lengths of SAT2 FISH, IdU, and YOYO-1 signals (in µm) were measured using ImageJ. Results are expressed as the % replicated SAT2 = [*Σ* (Length IdU tracts) / *Σ* (Length SAT2 DNA tracts)] x 100% or % Total replicated DNA = [*Σ* (Length IdU tracts) / *Σ* (Length of YOYO-1 tracts)] x 100%.

##### To assess fork stalling

Replication tracts in SAT2 were identified by the simultaneous detection of IdU, CldU, and SAT2 DNA using Genomic Vision’s EasyComb service. Images were acquired using the FiberVision scanner (Genomic Vision) and the fibers were manually measured using FiberStudio software (Genomic Vision). Analysis of fork symmetry was done by comparing the lengths of IdU and CldU incorporation into right- and left-moving sister forks. An asymmetric fork is defined as one where the ratio of the length of the right- and left-moving sister forks is <0.8 or >1.25.

